# A large phylogenetic tree for euphyllophytes

**DOI:** 10.64898/2026.01.06.695000

**Authors:** Tom Carruthers, William J. Baker, Wolf L. Eiserhardt, Félix Forest, Alexandre R. Zuntini, Jared T. Miller, Stephen A. Smith

## Abstract

**Premise:** Molecular datasets for estimating phylogenetic trees increasingly include more species and gene regions. Often trees are constructed using backbone phylogenies, subtrees, and other techniques to address the challenges of large dataset size. Currently, there is no established approach to integrate these rapidly expanding datasets.

**Methods:** We generated a phylogenetic tree (and 1,000 bootstrap trees) with divergence times that span euphyllophytes. To do this, we integrated taxonomically broad dated backbone phylogenies with species-level trees generated from phylogenetic analysis of individual clades. Datasets for species-level trees were assembled using *PyPHLAWD*.

**Results:** The resulting dated phylogenetic tree includes: 121,641 angiosperm species; 1,026 gymnosperms; and 5,603 ferns. This is the largest euphyllophyte phylogenetic tree constructed to date. Topological uncertainty spikes at the start of the Cretaceous and gradually increases during the Cenozoic. Uncertainty in age estimates gradually increases in the Cenozoic but increases dramatically in the most recent 5 Myrs.

**Discussion:** This dated phylogenetic framework can underpin evolutionary studies spanning euphyllophytes, and enable the integration of insights from the recent to the distant past. Our approach also enables the tree to be easily updated in the future to reflect future increases in data availability, and systematic and taxonomic advances within specific clades.

## INTRODUCTION

By representing relationships among taxa, phylogenetic trees underpin biological classification and our knowledge of the evolutionary processes that generate and maintain biological diversity. Phylogenetic trees with branch lengths proportional to time, or dated phylogenetic trees, are especially powerful for investigating evolutionary processes because they enable analyses of how changing biotic and abiotic environments shape the evolution of lineages through time.

The previous few decades have seen a profound increase in the quantity of molecular data from which phylogenetic trees can be estimated. This has transformed understanding of evolutionary relationships and underlying processes, from those underpinning major clades in land plants (e.g. Qui et al. 2006; 1KP initiative 2019) or animals (e.g. Adoutte et al. 2000; Schultz et al. 2023), to those underpinning relationships among more closely related species (Larson et al. 2020; Deanna et al. 2025). In some relatively small but charismatic major clades, such as birds, it is now possible to estimate phylogenetic trees that include over 95% of accepted species, sample 100s of genes for a large proportion of species, and are based on a stable taxonomic framework (McTavish et al. 2025).

This is not the case for land plants. With at least 400,000 species (Nic Lughadha et al. 2016), generating genomic-scale datasets that sample a significant proportion of species throughout the clade is very challenging. For example, the largest genomic-scale sequencing effort for angiosperms samples approximately 0.02% of all angiosperm species (Zuntini et al., 2024). Furthermore, even if dramatically more data were available, taxonomic uncertainty and computational/methodological limitations would still preclude the generation of robust phylogenetic trees for many clades (e.g. Carruthers et al. 2024; Zhang et al. 2025).

Despite these acute challenges, ongoing sequencing efforts are making major contributions. For example, the dataset in Zuntini et al. (2024) is unprecedented because it samples all flowering plant families and nearly two-thirds of all genera across 353 single- or low-copy nuclear gene regions. This provides a data-rich foundation for understanding relationships across the breadth of flowering plants, which is continually updated (Baker et al. 2022). Alongside broad-scale projects, a much larger number of studies with a narrower taxonomic focus continually generate data that is deposited in GenBank. Together, these data represent a much higher proportion of species than broad-scale sequencing projects, enabling the estimation of relationships between close relatives. For example, in 2018, GenBank data facilitated the construction of a phylogenetic tree for seed plants comprising approximately 80,000 species (Smith and Brown 2018). Although much of the data available at narrower scales is from Sanger sequencing (and thus samples relatively few loci), an increasing number of projects are generating densely sampled datasets underpinned by 100s of gene regions. Generating a single phylogenetic framework that integrates the strengths of broad and narrow-scale datasets can enable high-species-sampling evolutionary analyses anchored within a data-rich, broad-scale framework.

Here, we build on the approach set out in Smith and Brown (2018) to generate a dated euphyllophyte phylogenetic tree. This dated phylogenetic tree combines broad-scale molecular datasets such as that of Zuntini et al. (2024), with more densely sampled data for specific clades assembled from GenBank using *PyPHLAWD* (Smith and Walker, 2018). Using the dated phylogenetic tree from Zuntini et al. (2024), the Magallón et al. (2015) tree, which was used as a backbone in Smith and Brown (2018), is replaced. Importantly, the dated phylogenetic tree presented here accounts for topological and temporal uncertainty using bootstrapping. The approach is also designed to ensure that the resulting phylogenetic tree can be updated easily and regularly to incorporate the latest publicly available data. This includes datasets from clade-specific studies that sample many loci (including phylogenomic/transcriptomic datasets) and provide a more thorough account of distinctive evolutionary processes (Shah et al. 2021; Pérez-Escobar 2024; Schley et al. 2025) or taxonomic uncertainty/revisions (e.g., Muñoz-Rodríguez et al. 2019; Deanna et al. 2025) compared to the datasets assembled from GenBank. The breadth and species sampling of the phylogenetic trees generated by this project mean they will be useful to those interested in evolutionary questions spanning a wide range of taxonomic and temporal scales.

## MATERIALS AND METHODS

### Assembly of species-level datasets

We connected our taxonomy to both the World Checklist of Vascular Plants (WCVP) (Govaerts et al. 2021), and the National Center for Biotechnology Information (NCBI) (Schoch et al. 2020). To link these databases, we developed a relational framework for linking accepted species names from WCVP to all name associations in both NCBI and WCVP. Specifically, for each WCVP accepted name, we assembled a relational table that grouped together all relevant names, accepted names and synonyms, from both WCVP and NCBI. For every such taxon group, we tracked all corresponding NCBI identifiers (including those for accepted names, as well as synonyms and infraspecifics), ensuring that genetic sequence data would be accessible post-alignment. When conflict occurred between nomenclatures (i.e. NCBI accepted a name that WCVP treated as a synonym), we standardized the taxonomy according to WCVP and mapped the NCBI accepted name and identifier as a synonym to its WCVP counterpart. This ensured that taxon names retained WCVP relationships while retaining NCBI identifiers to source genetic data. As a last step we collapsed all infraspecific names to their parent species to simplify taxonomic units for phylogenetic inference, ensuring that all NCBI identifiers from infraspecific names and their synonyms were included.

All code was implemented in R v.4.3, using the packages data.table (1.17.6) (Barret et al. 2024), rgnparser(0.3.0) (Chamberlain et al. 2023), tidyverse(2.0.0) (Wickham 2023), and taxadb(0.2.1) (Boettiger et al. 2023). We first loaded the WCVP taxonomic backbone from a csv file obtained on December 2, 2024 and loaded the NCBI taxonomic backbone using taxadb::taxa_tbl(). The NCBI backbone was restricted to Viridiplantae by filtering the kingdom field, and names with ambiguous identifiers (e.g., “cf”, “x”, “sect”) were excluded to minimize uncertainty and exclude hybrids. For NCBI names, we selected key fields and parsed scientific names into canonical forms and authorship using rgnparser::gn_parse_tidy(). Names in NCBI were reformatted to indicate which names were synonymous with accepted names. Names mapping to multiple accepted names were excluded, given their ambiguous relationships and their minority representation (<5%) in NCBI. For WCVP’s taxonomic backbone, similar methods were used to construct relationships across accepted names and their synonyms, with one exception: accepted names that were subject to multiple name relationships were retained to preserve the name itself for downstream processing. ‘Unplaced’ names were removed from the relations entirely. Subsequently, any intraspecific information associated with names was removed.

A full join was performed on the name fields to align the two backbones, prioritizing WCVP names and backfilling NCBI names (including NCBI identifiers) where necessary. Due to conflicts between backbones and the simplification of accepted names to their parent taxon names, there were cases in which multiple NCBI identifiers existed for the same accepted name. These identifiers were retained and ordered, prioritizing WCVP’s version of the name for retrieving genetic data. These simplifications represent a key limitation when combining genetic databases with modern taxonomy to produce phylogenetic trees, potentially introducing inconsistencies relative to a manual approach.

We used this relational approach (rather than standard name matching) for the following reasons. First, a minority of names (<5%) map to multiple accepted taxa based solely on authorship, which presents challenges for linking to datasets that have non-standardized or missing authorities. Scanning through both databases, we were able to identify and remove any such names. Second, reconciling NCBI taxonomy with WCVP required accounting for differences in taxon concepts, not just names. In addition to the benefits of connectivity that our approach brings, the WCVP also has additional information about geography and characters that give a significant added benefit to datasets that forge a connection. The WCVP also continues to be updated and we will use updated versions as we construct new phylogenies.

### Phylogenetic inference for species-level datasets

We employed an approach based on that of Smith and Brown (2018), but modified in several respects, to construct the individual species-level phylogenetic trees. First, before downloading data from NCBI, we filtered out sequences that contained the terms "Patent", "unplaced", "retrotransposon", "scaffold", and "like element" in any field. This significantly shrinks the available data for phylogenetic analysis and makes subsequent analyses simpler. This is also an alternative to downloading the entire “pln” dataset from the NCBI ftp. Additionally, a custom Python script was used to query the PyPHLAWD NCBI-style SQLite taxonomy database, construct a rooted tree for a user-specified taxon (optionally restricted to a provided list of taxa), and export the resulting tree in Newick format using a lightweight node-based tree library. The script then traversed the tree to identify internal clades with fewer than 15,000 descendant tips and, for each such clade, reported a suitable outgroup defined as a sister subtree with at least 10 leaves that does not contain any of the focal clade’s descendants. These clades, with fewer than 15,000 taxa, helped us identify units that we could analyse independently to date and graft to the backbone later. This size was determined to be sufficient to reduce the number of individual trees while still enabling large-scale analyses without excessive analysis time.

We used an automated approach to assembling the final datasets based on the clusters constructed by PyPHLAWD. Specifically, phylogenetically informed clusters were included in the final dataset when all their taxa were contained within a single node of the directory-derived tree and provided that the cluster either represented at least 20% of the taxa in that node (cluster_prop = 0.2) or included at least 20 taxa in total (smallest_cluster = 20). Only clusters with ≥4 taxa that met these coverage/size criteria at an internal node or the root were included in the final datasets (with smaller clusters optionally included via the “includetrivial” setting). Upstream and optional filters such as a minimum sequence length of 450 bp (smallest_size = 450) and tree-trimming cutoffs (relcut = 0.25, abscut = 0.7, abscutint = 1.5) did not alter this basic cluster inclusion rule. As in the original Smith and Brown (2018) publication, a topological constraint tree was generated by querying a local SQLite instance of the NCBI taxonomy restricted to the sampled taxa. To prevent the enforcement of ambiguous or redundant constraints, the script collapsed all unifurcating (single-child) nodes. It merged lineages designated as ‘incertae sedis’ or ‘unidentified’ directly into their parent nodes. Unlike Smith and Brown (2018), this constraint topology was then refined by reconciling it with a reference PAFTOL phylogeny. To do this, leaf labels were mapped to NCBI taxon IDs (restricted to Viridiplantae), and binary splits observed in the reference backbone were enforced in the constraint tree by inserting internal nodes where daughter clades appeared as sisters, thereby resolving polytomies consistent with the reference while preserving the original leaf labels.

Before conducting a clade-specific analysis, we augmented the existing concatenated matrix with outgroup sequences using a custom BLAST-based pipeline. For a specified NCBI taxon, we extracted non-genome sequences from a precompiled GenBank database and filtered for regions with >100 ungapped base pairs. We queried these sequences against gene partitions defined by the original matrix using BLASTN (perc_identity = 20, E-value = 1×10−9, multi-threaded), retaining the best-scoring hit for each taxon–gene combination. Taxa meeting a user-defined locus threshold (minoverlap; default = 1) were selected for inclusion, up to a maximum limit (maxtaxa; default = 1). The selected sequences were aligned to their respective gene regions using MAFFT and concatenated with pxcat to generate an updated supermatrix, partition file, and summary table.

Using RAxML version 8.2.12 (Stamatakis 2014), we conducted maximum-likelihood runs with the constrained relationships. These were done with the GTR+CAT model using the concatenated matrix. Specifically, we ran with the -f a option and 100 rapid bootstraps. We initially used the -U option for autoMRE bootstrapping. However, this caused some challenges for branch-length estimates in the bootstrap runs. Therefore, we reran those branch-length estimates under the GTR+Γ model and rerooted using the outgroups identified above. We conducted 108 clade-specific runs.

### Divergence time estimation for backbone tree spanning euphyllophytes

To incorporate sufficient phylogenetic breadth and allow for the integration of species-level trees spanning euphyllophytes, we combined several datasets to generate dated backbone-trees (Fig. 1a). First, we utilised the transcriptomic dataset of Pelosi et al. (2022). This dataset (specifically their “SCO60” dataset) samples 2,800 loci for 244 fern species, and also includes samples for gymnosperms, angiosperms, lycophytes, and bryophytes. It therefore provides a relatively comprehensive backbone for ferns whilst including sufficient outgroups such that it can be integrated with other clades (i.e. gymnosperms, angiosperms, see below.)

**Figure 1.**
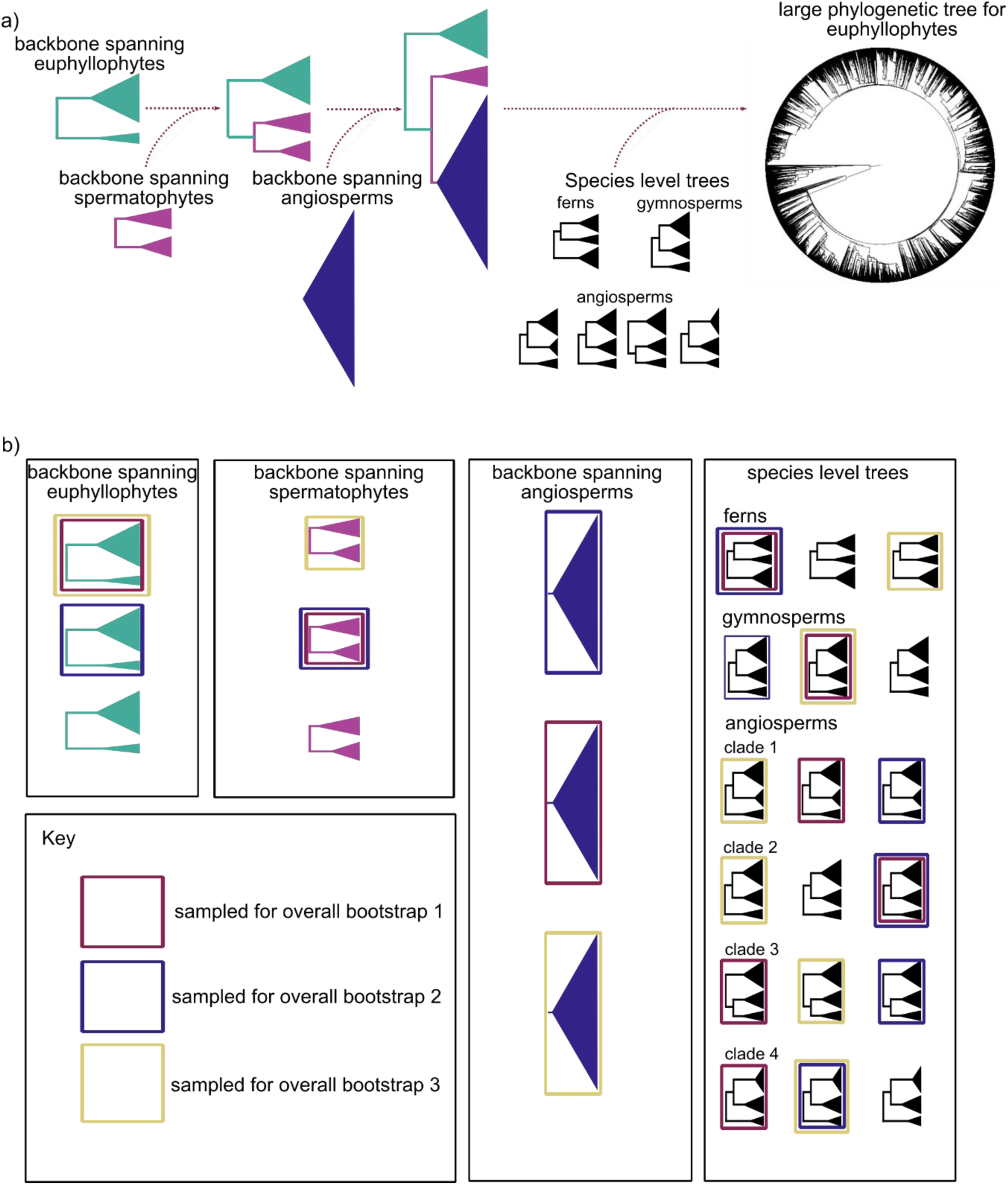
Overview of the assembly of large phylogenetic trees for euphyllophytes. a) Illustration of how the different backbone trees and species-level trees correspond to each other and are combined to generate the overall trees. b) Illustration of the bootstrap sampling procedure. Different pools of trees are shown in each black outlined box. In this figure, there are only three within each pool, but 100 are generated in this study. Trees from these different pools are randomly sampled to generate an overall bootstrap tree. The trees sampled to generate each overall bootstrap tree are surrounded by a coloured box, as highlighted in the key. In this study, we repeat the procedure of randomly sampling trees from each pool 1000 times, rather than the three times displayed in this figure.

We re-ran divergence time analyses on this dataset in order to generate a dated backbone tree that was methodologically consistent with the other dated backbone trees used in this study. In contrast to Pelosi et al. (2022), we therefore removed the *P. patens* outgroup prior to divergence time estimation; adjusted some fossil calibrations (see below); and ranked loci solely on the basis of topological congruence with the species tree. To achieve this, the methodological steps outlined below were followed.

First, loci were ranked using SortaDate (Smith et al., 2018) based on the proportion of congruent bipartitions between each gene tree and the species tree. Aligned sequences (from Pelosi et al. (2022)) for any locus in which the gene tree had at least 75% of congruent bipartitions with the species tree were concatenated (101 loci). This concatenated alignment was then used to re-estimate molecular branch lengths in RAxML, with the topology constrained to the SCO60 topology of Pelosi et al. (2022), and a GTRCAT model that was partitioned by each locus. Maximum likelihood branch lengths and 100 bootstrap branch lengths were estimated.

The maximum likelihood and bootstrap trees were then rooted with *Physcomitrella patens*. *P. patens* and *Selaginella moellendorffii* were then pruned from the maximum likelihood tree and each bootstrap tree, before the trees were dated in *treePL* (Smith and O’Meara 2012). Fossil calibrations were similar to those of Pelosi et al. (2022) (Table S1), with the following exceptions. 1) A fossil calibration at the crown node of land plants (based on a Cryptospore fossil) was no longer necessary. 2) The euphyllophyte crown node (corresponding to the root of the pruned trees), was constrained with a minimum of 388 Ma and a maximum of 423 Ma, as in Stull et al. (2021). The minimum corresponds to *Rellimia* (Dannenhoffer and Bonamo 1989) – the earliest fossil evidence for Progymnosperms, whilst the maximum corresponds to *Baragwanathia* (Garrat 1978), the oldest evidence of crown vascular plants. 3) The crown node of seed plants was constrained with a minimum of 318 and a maximum of 366.8 Ma. The maximum corresponds to *Elkinsia polymorpha* (Rothwell et al. 1989), the earliest known fossil seed plant lineage, and we follow Stull et al. (2021)’s interpretation that crown group seed plants are unlikely to significantly predate this age. The minimum corresponds to Cordaitales fossils in the Carboniferous (Namurian) (Phillips 1980; Taylor et al. 2009). 4) A maximum constraint for the flowering plant crown node of 154 Ma was used, consistent with Zuntini et al. (2024). 5) The Kuylisporites and Glomerisporites fossils applied at the crown nodes of *Alsophila* + *Cyathea* and *Azolla* were moved to the stem nodes of their respective clades following Collinson (2001) and Batten et al (1998). Further, except where stated above, all maximum age constraints from Pelosi et el. (2022) were removed. In that study, those maximum constraints correspond to the lower bound of the stratigraphic interval from which the fossil was sampled. Although this corresponds to the maximum possible age of the fossil, it does not necessarily correspond to the maximum possible age of the clade.

Cross-validation, following an initial priming step, was performed in *treePL* on the maximum likelihood tree and the calibrations described above. Smoothing values between 10,000 and 0.001 were trialled, with a smoothing value of 1 yielding the lowest χ². This smoothing value was used to estimate divergence times on the maximum likelihood tree and all bootstrap trees.

### Divergence time estimation for backbone tree spanning spermatophytes

The transcriptomic dataset of Ran et al. (2018), which samples all 13 gymnosperm families (plus 13 angiosperm species and 3 fern species) was included to provide a backbone for gymnosperms (Fig. 1a). Similarly to our analysis of the Pelosi et al. (2022) dataset, alignments for loci with gene trees that were most congruent with the species tree topology from Ran et al. (2022) (all species trees in that study are topologically identical) were concatenated. Molecular branch lengths were then re-estimated on the species tree topology from Ran et al. (2022) in RAxML with a GTRCAT model partitioned by locus. A maximum likelihood tree and 100 bootstrap trees were estimated.

The maximum likelihood and bootstrap trees were rooted with ferns as the outgroup, and then the outgroup was pruned. This leaves a rooted tree comprising gymnosperm and angiosperm samples i.e. a tree spanning spermatophytes. Divergence time estimation was then undertaken in *treePL*. Fossil calibrations are summarised in Table S2. These are primarily based on those used in Stull et al. (2021), but include some additional calibrations within angiosperms that were used by Zuntini et al. (2024). All calibrations were implemented as minimum age constraints, except a 154 Ma maximum constraint at the crown node of angiosperms, as used in Zuntini et al. (2024). Further, in the maximum likelihood tree, the root node (i.e. the spermatophyte crown node) was constrained to be the same age as the spermatophyte crown node in the dated maximum likelihood backbone tree spanning euphyllophytes. Meanwhile, for each bootstrap tree, the root node was constrained to be the same age as the spermatophyte crown node in a randomly sampled bootstrap from the bootstrap backbone trees spanning euphyllophytes.

As before, cross-validation and priming were performed on the maximum likelihood tree. A smoothing value of 10 had the lowest χ^2^ value and was used when dating the maximum likelihood tree and all bootstrap trees.

### Divergence time estimation for angiosperm backbone tree

The backbone tree for angiosperms is based on the tree from Zuntini et al. (2024) (Fig. 1a), in which molecular branch lengths were estimated from a concatenated alignment of the 25 loci with gene trees that were most congruent with the species tree topology. In that study, molecular branch lengths were estimated from these 25 loci using IQTREE 2 (Minh et al. 2020), with the topology constrained to that estimated in their main ASTRAL (Mirarab et al., 2014) analysis. We used the same analytical setup to generate a further 100 bootstrap trees with molecular branch lengths.

The maximum likelihood tree from Zuntini et al. (2024) plus the 100 bootstrap trees were dated in *treePL.* The same fossil calibrations as Zuntini et al. (2024) were used, except that the root node was constrained to be identical to the age estimate of the angiosperm crown node from the backbone tree spanning spermatophytes. However, in the maximum likelihood tree and all bootstrap trees, this node was estimated to be 154 Ma. Therefore, in practice, fossil calibrations were identical to Zuntini et al. (2024). A smoothing value of 10 was used, as per Zuntini et al. (2024).

### Divergence time estimation in species-level datasets

For each of the 108 species-level datasets (106 for angiosperms, 1 for gymnosperms, and 1 for ferns) divergence times were estimated on the maximum likelihood tree and the 100 bootstrap trees. Age constraints for the species-level trees were based on divergence time estimates in the relevant backbone tree, i.e. angiosperm species-level trees had age constraints based on the angiosperm backbone trees, gymnosperm species-level trees had age constraints based on the spermatophyte backbone trees (which is most densely sampled in gymnosperms), and the fern species-level trees had age constraints based on the euphyllophyte backbone trees (which is most densely sampled in ferns) (Fig. 1a).

Specifically, for each species-level tree, nodes that overlapped with nodes in the relevant backbone tree (e.g., if both the species-level tree and backbone tree include the crown node for a genus) were constrained to have the same age as the node in the backbone tree. If the backbone tree samples one species for a genus, and there are multiple species for that genus in the species-level tree, the age of the node immediately ancestral to the tip for that genus in the backbone tree is used as a maximum age constraint for the crown node of the genus in the species-level tree.

In some cases, a species-level tree may sample species that are outside the respective clade in the backbone tree for which the species-level dataset was generated. A further step is implemented in such cases where the root node of such species-level trees has a maximum age constraint equal to the stem node age of the relevant clade in the backbone tree. In a handful of cases, the species-level tree corresponds to a single tip in the backbone tree. In such cases, the maximum age constraint for the root node of the species-level tree is equal to the age of the node that is immediately ancestral to the relevant tip in the backbone tree.

The backbone tree from Zuntini et al. (2024) contains numerous non-monophyletic taxa (in particular genera, but some families). These clades are often overlooked when setting constraints, or if a single outlier tip causes non-monophyly, it is ignored. In some cases, there are also incongruences between the species-level trees and backbone trees – for example, where the NCBI taxonomy has been favoured over the Zuntini et al. (2024) topology when setting topological constraints for the species-level trees. Time constraints cannot be set for such clades within the species-level trees.

The same process was followed when generating node age constraints for the maximum likelihood species-level trees and bootstrap species-level trees. For the maximum likelihood trees, the age constraints are derived from the relevant maximum likelihood backbone tree. Age constraints for species-level bootstrap trees are derived from a randomly sampled backbone bootstrap tree (more details below). Age estimates for each species-level tree were estimated in *treePL* with a smoothing value of 10, following an initial priming step.

### Integrating trees to generate a single maximum likelihood tree and 1,000 bootstrap trees

To generate a single tree from the maximum likelihood trees, the backbone trees were grafted together (Fig. 1a). Specifically, the spermatophyte backbone tree was grafted to the euphyllophyte backbone tree at the spermatophyte crown node. Then, the angiosperm backbone tree was grafted to the angiosperm crown node in the combined euphyllophyte and spermatophyte backbone tree (Fig. 1a). Species-level trees were then grafted onto the crown nodes of their clade in the relevant backbone tree (Fig. 1a). Where species-level trees had a maximum constraint at the root node derived from the stem node age of the relevant clade in the backbone tree (see above), species-level trees were instead grafted onto the stem lineage of their relevant clade at a height that corresponded to the root node age estimate in the species-level tree. Further, species-level trees that corresponded to single tip in the backbone tree were grafted onto the branch leading to that tip at a height that corresponded to the root node age estimate in the species-level tree. Subsequently, all tips in the backbone trees that corresponded to clades or tips to which species-level trees had been grafted were removed. An undated tree was also generated using the undated trees (following rooting and removal of outgroups) and the same grafting procedure.

A similar process is followed for the bootstrap trees, except that trees are grafted to the randomly sampled bootstraps to which they are constrained. To clarify, the overall procedure for randomly sampling bootstrap trees involves randomly selecting one tree from each of 100 bootstrap trees from the euphyllophyte, spermatophyte, and angiosperm backbones, as well as the 108 species-level datasets (Fig. 1b). In the current study, we repeated this process 1000 times to generate a total of 1000 combined dated and undated bootstraps.

### Including species with no molecular data

Alongside the phylogenetic trees described above, where every species has molecular data, we present a further phylogenetic tree where species without molecular data are added. Specifically, additional species for genera represented in the phylogenetic tree described above (as per WCVP), are added as unresolved tips in their respective genus.

### Updating the tree and generating further bootstraps

An updated set of trees reflecting updates in the availability of species-level data will be generated and deposited every six months at: https://doi.org/10.17605/OSF.IO/9TBHA. All trees generated as part of the study are deposited here: https://doi.org/10.17605/OSF.IO/9TBHA, and scripts to generate further bootstraps are available at: https://doi.org/10.5281/zenodo.17968787. An R package is also available (https://github.com/jtmiller28/EuphyllophyteTreeRetriever) to enable easy access to the trees – specifically the download of bootstraps spanning the entire of Euphyllophytes, as well as bootstraps for user defined clades.

Users may wish to substitute phylogenetic trees underpinned by extensive clade specific taxonomic/systematic studies into the overall tree presented here. Such trees are likely to more accurately reflect taxonomic knowledge within specific clades compared to our automated approach. We present a set of R functions (available at: https://doi.org/10.5281/zenodo.17968787) that enable users to do this easily. Specifically, these functions take an undated phylogenetic tree for a specific clade (or set of bootstrap trees), generate time constraints that are consistent with the overall tree, date the clade specific tree(s), and graft them into the overall tree. These functions are highly flexible, enabling the replacement of clades spanning multiple orders, or clades comprising groups of species within a single genus. As an example, we present an updated version of our dated phylogenetic tree where Ericales has been replaced with the Ericales phylogenetic tree from Carruthers et al. (2024) (available here: https://doi.org/10.5281/zenodo.17968787).

## RESULTS

### Data assembly

The number of species in the molecular datasets from which species-level trees were estimated varied from 4 to almost 12,000 (mean = 1196.426 species), whilst the number of genes included in each dataset varied from 1 to 296 (mean = 32.4537; Table S3). Typically, datasets with more species included a larger number of genes (Fig. 2a). The mean sequence length varied considerably between datasets from a minimum of 551 for Picraminales to 27577.72 for Icacinales (Fig. 2b, Table S3). Further statistics on the assembled molecular datasets are in Table S3.

**Figure 2.**
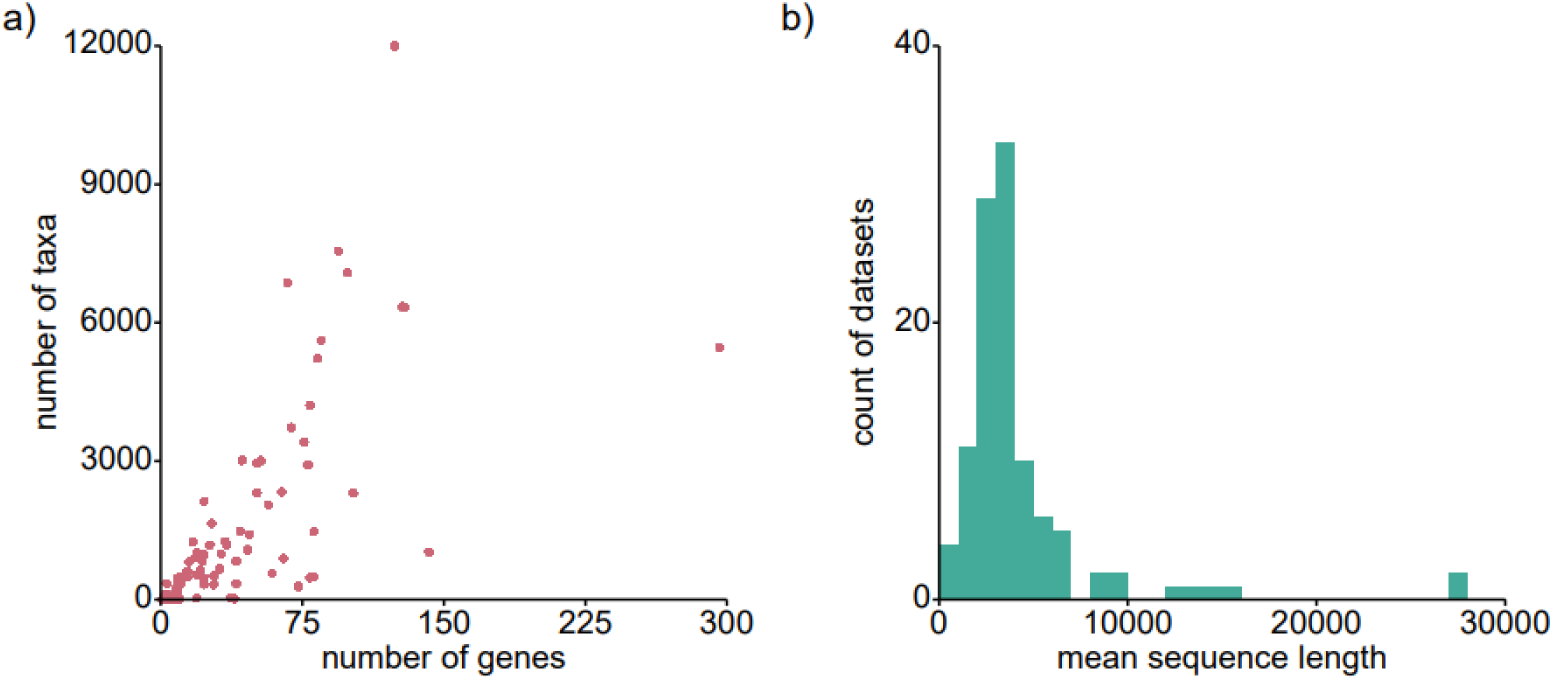
Summary of assembled molecular datasets from which species-level trees were estimatred. a) number of genes in each dataset plotted against the number of taxa in the dataset. b) Mean sequence lengths across all the species-level datasets.

### Final combined phylogenetic tree

The final combined phylogenetic tree for euphyllophytes has 128,270 species. Of these, 5,603 are ferns (∼46% of accepted fern species), 1,026 are gymnosperms (∼84% of accepted gymnosperm species), and 121,641 are angiosperms (∼31% of accepted angiosperm species). Within angiosperms, the most densely sampled clades are primarily temperate such as Asteliaceae, Restionaceae, or Goodeniaceae, whilst predominantly tropical clades, including large clades such as Pandanales, Dilleniales, or Acanthaceae, are comparatively incompletely sampled (Table S4). 53 species from the angiosperm backbone tree were retained in the final phylogenetic tree, with all other species being removed when the species-level trees were grafted.

Bootstrap values are relatively high for the most deeply nested nodes in the phylogenetic tree. This is likely due to a high proportion of nodes in this part of the tree that are constrained to the topology of the backbone tree (Fig.3a). Bootstrap values than show a pronounced drop in the Late Jurassic and Early Cretaceous (Fig.3a). This corresponds to the origin of Polypodiales in ferns, as well as major angiosperm clades such as Asparagales, Ericales, and Ranunculales. Topological uncertainty then gradually decreases during the second half of the Cretaceous, before increasing gradually in the Paleogene, and more rapidly in the Neogene and Quaternary. Bootstrap values for specific angiosperm clades (calculated as a mean across all nodes for each species-level tree) were lowest in Blandfordiaceae, Juncaceae, and Fagales, and highest in Martyniaceae, Stilbaceae, and Thurniaceae. As expected, bootstrap values are higher in clades for which a higher proportion of nodes overlap with nodes included in Zuntini et al. (2024) (Fig.3b).

**Figure 3.**
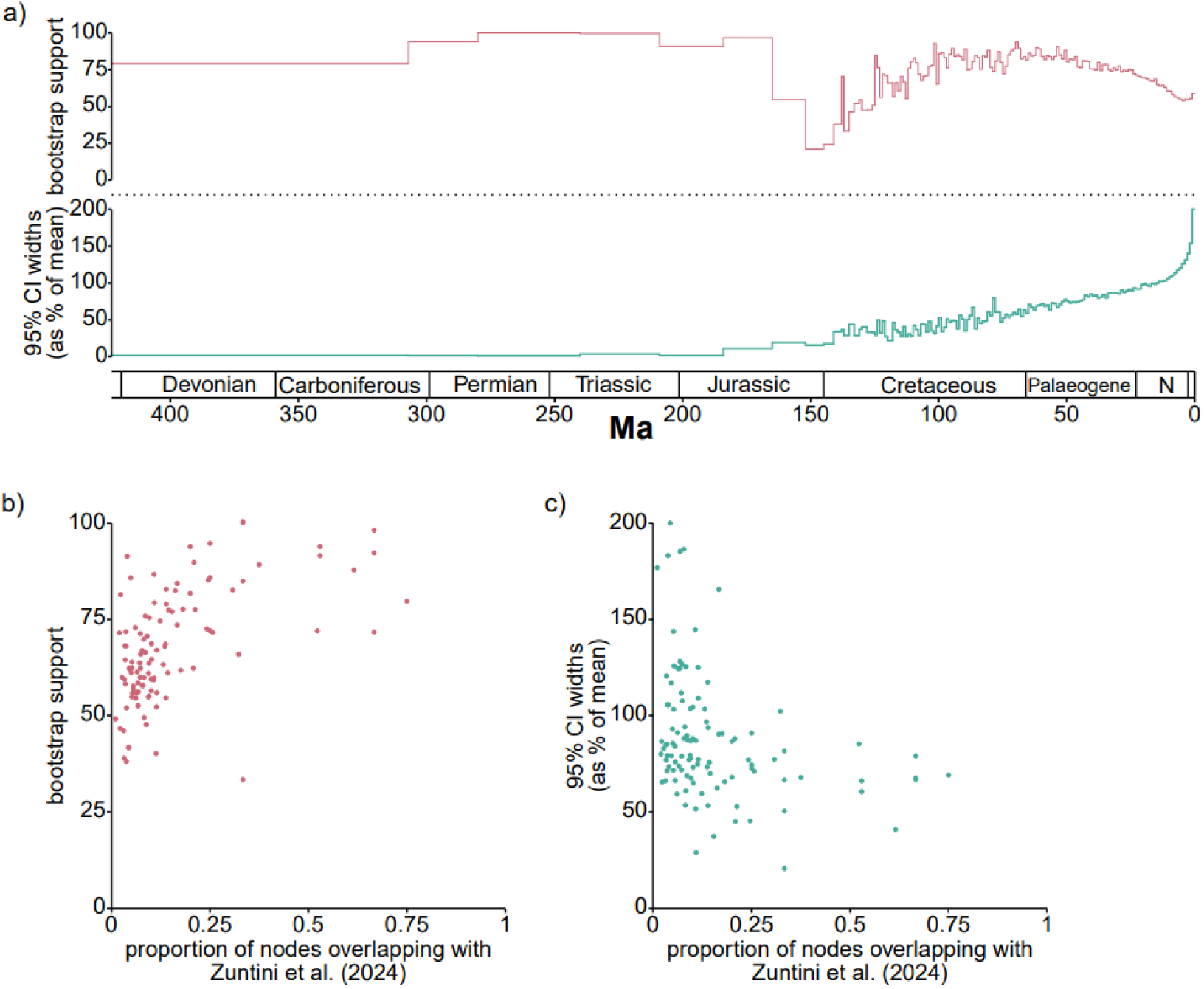
Summary of topological uncertainty and uncertainty in age estimates. a) Uncertainty in topology and age estimates through time. Values are calculated as a mean across all nodes within 1Myr time intervals. If there are fewer than 5 nodes in the time interval, the time interval is extended backward in time until it includes 5 nodes. For uncertainty in age estimates i.e. 95% CI widths (as % of mean), the mean refers to the mean age estimate for the node across all bootstraps. b) The mean bootstrap support for angiosperm species-level trees plotted against the proportion of nodes that overlap with Zuntini et al. (2024). c) The mean 95% CI widths (as % of mean) plotted against plotted the proportion of nodes that overlap with Zuntini et al. (2024).

### Divergence time estimates

At the broadest scale, the age estimate for the euphyllophyte crown node was 423 Ma, spermatophytes was 318 Ma, gymnosperms was 306.2 Ma, and angiosperms was 154 Ma. These age estimates are consistent across all bootstraps and correspond directly to the maximum age constraints applied at the euphyllophyte and angiosperm crown nodes and the minimum age constraints applied at the spermatophyte and gymnosperm crown nodes. This implies that these calibrations have a significant impact on age estimates. Other users may want to experiment further with these calibrations, which is straightforward to do with the code provided on Zenodo. These age estimates are broadly consistent with the fossil record, and many previously published molecular-based age estimates (Magallón et al. 2013; Stull et al. 2021; Zuntini et al. 2024). At narrower scales, age estimates are, unsurprisingly, broadly consistent with the backbone trees on which this study is based. Nonetheless, by including many more nodes, the dated phylogenetic framework presented here significantly extends insights into the timing of plant evolution compared to previously published dated phylogenetic trees.

As with topological uncertainty, uncertainty in age estimates is low on deep branches in the phylogenetic tree (Fig.3a). Unlike topological uncertainty, there is no significant increase in uncertainty in age estimates at the beginning of the Cretaceous. Uncertainty in age estimates then increases gradually from the second half of the Cretaceous (similarly to topological uncertainty), before spiking during the Quaternary. As with topological uncertainty, uncertainty in age estimates tends to be higher in angiosperm clades in which a lower proportion of nodes are included in Zuntini et al. (2024) (Fig.3c, Table S4).

## DISCUSSION

We present a dated phylogenetic tree for euphyllophytes that includes approximately one-third of all species in this clade, and samples evolutionary branching events spanning tens of thousands to over 400 million years ago. Deeper relationships within the tree are informed by genomic-scale datasets, including that of Zuntini et al. (2024), which samples nearly 2/3 of all flowering plant genera. Meanwhile, shallower branching events are estimated from the increasingly large quantities of data available in public repositories. This phylogenetic tree provides a foundation for studying plant evolution across a much broader range of temporal and taxonomic scales than has been possible previously. By generating 1,000 bootstrap trees, we also quantify topological uncertainty and uncertainty in age estimates. Users of this phylogenetic framework can therefore perform analyses that explicitly account for uncertainty.

Our results relating to uncertainty are consistent with previous studies and highlight areas where the phylogenetic tree can be improved in the future. For example, the increase in topological uncertainty in the Early Cretaceous is consistent with the well-established phenomenon that the early evolution of major angiosperm clades is associated with topological uncertainty and conflict among gene regions (Larson et al. 2020; Yang et al. 2023; Smith et al. 2025). Likewise, our finding that high topological uncertainty during early angiosperm evolution is not associated with uncertainty in age estimates is consistent with previous findings in which molecular divergence time estimates for early angiosperm nodes are primarily sensitive (when using existing molecular clock methods) to the maximum constraint at the angiosperm crown node, regardless of the topology (Magallon et al. 2015, Zuntini et al. 2024). The marked increase in uncertainty in the last 5Ma, especially for age estimates, likely corresponds to the fact that during this timeframe, the number of nodes that overlap with Zuntini et al. (2024) decreases markedly. The tree we present may therefore be improved in the future by incorporating species-level genomic-scale datasets underpinned by extensive clade-specific study.

We do, however, note that some caution should be taken when interpreting the uncertainty that we present. First, topological uncertainty may be underestimated on deeper branches in the tree, given that the backbone trees have the same topology across all bootstraps. This is unlikely to be especially important for gymnosperms and ferns (given the vast majority of these nodes have high support in their respective studies) (Pelosi et al. 2022; Ran et al. 2022), but may be more important in angiosperms. More effectively incorporating this uncertainty in the future may lead to even greater topological uncertainty in the Early Cretaceous. Alongside this, the confidence intervals we present for age estimates should be interpreted in the context of the limited set of assumptions about molecular rate variation employed in this study. Age estimates are very sensitive to such assumptions (Carruthers et al. 2020; Smith and Beaulieu 2024), and equally plausible age estimates would likely be obtained for many nodes that lie outside the confidence intervals presented here.

The approach presented here can enable the generation of updated trees to be readily accomplished in the future. These might include the latest data available on GenBank, phylogenetic trees, and datasets underpinned by extensive study for specific clades, or backbone trees that sample more species or more effectively incorporate uncertainty. This provides a basis for producing regularly updated phylogenetic trees to study plant evolution, reflecting the latest available data, taxonomic treatments, and analytical methods.

## Supporting information

Supplementary Table 1; Supplementary Tabe 2

## ACKNOWLEDGEMENTS

The authors acknowledge the following grant: NSF DEB 2325835.

## AUTHOR CONTRIBUTIONS

TC, SAS, JTM wrote the code and performed the analyses. TC wrote the manuscript. All authors contributed to the manuscript.

## DATA AVAILABILITY

Code used to estimate the phylogenetic trees and functions to replace parts of phylogenetic trees with user specified trees is available here: https://doi.org/10.5281/zenodo.17968787. All trees are available at: https://doi.org/10.17605/OSF.IO/9TBHA. An associated R package to enable easy retrieval of these trees is available at: https://github.com/jtmiller28/EuphyllophyteTreeRetriever.

