## Supplementary Table 1; Supplementary Tabe 2 for "A large phylogenetic tree for euphyllophytes": Supplementary_Tables.docx

**Table S1.** Fossil calibrations used in backbone tree spanning euphyllophytes.

| **Fossil** | **Node** | **Age (Ma)** | **Reference** |
| --- | --- | --- | --- |
| *Remillia* | Euphyllophyte Crown | 388 (Min) | Dannenhoffer and Bonamo 1989 |
| *Baragwanathia* | Euphyllophyte Crown | 423 (Max) | Garrat 1978 |
| Cordiatales | Spermatophyte Crown | 318 (Min) | Phillips 1980; Taylor et al. 2009 |
| *Elkinsia polymorpha* | Spermatophyte Crown | 366.8 (Max) | Rothwell et al. 1989 |
| Secondary Constraint | Angiosperm Crown Node | 154 (Max) | Zuntini et al. 2024 |
| Archaeocalamites | Equisetales stem node | 323.2 (Min) | Bateman 1991 |
| Grammatopteris | Osmundales stem node | 299.0 (Min) | Skog 2001 |
| Szea sinensis | Gleicheniaceae stem node | 270.6 (Min) | Yao & Taylor 1988 |
| Stachypteris | *Lygodium* stem node | 167.7 (Min) | Van Konijnenburg-Van Cittert 1981, Wikstrom et al. 2002 |
| Regnellites | Marsileaceae stem node | 140.2 (Min) | Yamada & Kato 2002 |
| Kuylisporites | *Alsophila* + *Cyathea* stem node | 93.5 (Min) | Collinson 2001 |
| Glomerisporites | *Azolla* stem node | 83.5 (Min) | Batten et al. 1998 |
| *Acrostichum* | *Acrostichum* | 65.5 (Min) | Bonde & Kumaran 2002 |
| *Woodwardia gravida* | *Woodwardia* stem node | 55.8 (Min) | Wilf et al 1998; Collinson 2001 |
| *Makotopteris princetonensis* | *Athyrioids* stem node | 37.2 (Min) | Stockey et al. 1999 |
| *Protodrynaira sp.* | Polypodiaceae stem node | 33.9 (Min) | Vikulin and Bobrov 1987 |
| *Polypodium radonii* | Polypodium stem node | 25.6 (Min) | Kvacek 2001 |
| *Coniopteris lunzenis* | Polypodiales + Cyathaeales stem node | 228 (Min) | Samylina 1976, Dobruskina 1994, Schneider and Kenrick 2001, Schneider et al. 2004 |

**Table S2.** Fossil Calibrations in backbone tree spanning spermatophytes.

| **Fossil** | **Node** | **Age (Ma)** | **Reference** |
| --- | --- | --- | --- |
| Secondary Constraint | Spermatophyte crown | Node age estimated in backbone tree spanning Euphyllophytes |  |
| *Cordaixylon iowensis* | Acrogymnospermae crown | 306.2 (Min) | Trivett et al. 1992; Peppers 1996 |
| *Araucarities rudicula* | Araucariaceae + Podocarpaceae crown | 213.0 (Min) | Axsmith and Ash 2006 |
| *Austrohamia minuta* | Cupressaceae crown | 182.7 (Min) | Escapa and Bodnar 2017 |
| *Fokienia ravenscragensis* | Fokienia + Platycladus crown | 61.1 (Min) | McIver et al. 1990, 1992 |
| *Metasequoia* | Sequoioideae crown | 93.9 (Min) | LePage et al. 2005 |
| *Crossozamia* | Cycadales crown | 270.0 (Min) | Gao and Thomas 1989a,b |
| *Spermopteris* | Cycadales stem | 298.9 (Min) | Mamay 1976 |
| *Cratonia cotyledon* | Gnetaceae + Welwitschiaceae crown | 114.0 (Min) | Rydin et al. 2002 |
| *Ephedra drewriensis* | *Ephedra* stem | 125.0 (Min) | Rydin et al. 2006 |
| Polyplicate pollen | Gnetales stem | 265.1 (Min) | Wilson 1962 |
| *Eathiestrobus mackenziei* | Pinaceae crown | 151.1 (Min) | Rothwell et al. 2012 |
| *Picea burtonii* | *Picea* stem | 132.9 (Min) | Klymiuk and Stokey 2012 |
| *Palaeotaxus redivia* | Taxaceae stem | 197.0 (Min) | Florin 1958 |
| Secondary Constraint | Angiosperm Crown Node | 154 (Max) | Zuntini et al. 2024 |
| *Montsechia vidalii* | Ceratophyllum stem | 127.2 (Min) | Zuntini et al. 2024 |
| *Changii indicum* | Ehrhartoideae stem | 66.0 (Min) | Zuntini et al. 2024 |
| *Teixeiraea lusitanica* | Ranunculales stem | 110.8 (Min) | Zuntini et al. 2024 |
| *Hyrcantha decussata* | Eudicots stem | 125 (Min) | Zuntini et al. 2024 |
| *Dressiantha bicarpellata* | Brassicales stem | 86.3 (Min) | Zuntini et al. 2024 |
